# Brain organoid-on-a-chip to create multiple domains in forebrain organoids

**DOI:** 10.1101/2023.09.18.558278

**Authors:** Yuan-Chen Tsai, Hajime Ozaki, Ango Morikawa, Kaori Shiraiwa, Andy Prosvey Pin, Aya Galal Salem, Kenneth Akady Phommahasay, Bret Kiyoshi Sugita, Christine Hein Vu, Saba Mamoun Hammad, Ken-ichiro Kamei, Momoko Watanabe

## Abstract

Brain organoids are three-dimensionally reconstructed brain tissue derived from pluripotent stem cells in vitro. 3D tissue cultures have opened new avenues for exploring development and disease modeling. However, some physiological conditions, including signaling gradients in 3D cultures, have not yet been easily achieved. Here, we introduce Brain Organoid-on-a-Chip platforms that generate signaling gradients that in turn enable the induction of topographic forebrain organoids. This creates a more continuous spectrum of brain regions and provides a more complete mimic of the human brain for evaluating neurodevelopment and disease in unprecedented detail.

---

Brain organoids are self-organized three-dimensional structures derived from pluripotent stem cells (PSCs) that recapitulate many aspects of tissue structure *in vivo*.^1–3^ Organoid technology has tremendous potential for the study of human neurodevelopment and disease.^2,4^ However, many physiological conditions have not been achieved *in vitro*. For example, a localized group of brain cells during embryogenesis acts as a “signaling center” and creates gradients of secreted instructive signaling molecules to induce diverse domains and cell types in a concentration-dependent manner.^5^ This process is crucial for the formation of distinct brain regions. Currently, region-specific organoids are generated with a single-dose bath application of signaling molecules, which is beneficial for creating a single region but lacks gradients and thus cannot form multiple spatially organized regions.^4,6–8^ In order to create structures containing multiple regions using current approaches, distinct regions must be assembled separately despite the fact that these regions normally develop together *in vivo*.^9–13^ Methods to spontaneously make organoids containing multiple distinct brain regions lack sufficient reproducibility to enable well-controlled experiments to increase understanding of human brain development.^14^ Thus, the field needs a more reproducible method to generate forebrain organoids that reflect the structural complexity of the human brain.

To overcome these limitations, Organ(s)-on-a-Chip (OoC) platforms (also known as microphysiological systems) provide an unprecedented opportunity to study human pathophysiology as well as developmental processes *in vitro* for disease modelling and drug discovery.^15,16^ Most OoC platforms are based on microfluidic technology, which enables a broad range of biological applications by regulating environmental cues, including concentration gradients of soluble factors, 3D geometry and extracellular scaffolds, and perfusion.^17–19^ Microengineered devices have been used to generate brain-region-specific organoids^20–22^; however, those featuring multiple domains have remained elusive. This limitation stems from the challenge of generating concentration gradients of multiple signaling molecules for a single brain organoid. Although numerous microfluidic devices have been developed to establish concentration gradients for cellular stimulation, the perfusion flow frequently induces shear stress that damages cells.^23,24^ Therefore, there is a pressing need for devices that can apply concentration gradients to cultured organoids without significant shear stresses, thereby facilitating the creation of brain organoids with multiple domains.

Here, we developed Brain-Organoid-on-a-Chip (BOoC) to obtain topographically organized forebrain organoids by mimicking extracellular concentration gradients during development. First, we established a multi-layered microfluidic device to generate four kinds of concentration gradients in an organoid culture chamber without high shear stress. Second, we embedded a forebrain organoid into the chamber to be exposed to an extracellular smoothened agonist (SAG) concentration gradient. The resulting organoid exhibited topographically organized domains, including cortical, lateral, and/or medial ganglionic eminence-like regions in one organoid, whereas forebrain organoids with a bath application of SAG showed stochastic formation of domains with different regional identities or a single domain when a high concentration is used. Together, we demonstrate proof-of-concept that topographically organized forebrain organoids can be created using BOoC technology.

## Results

### Device design and fabrication

To generate topographically organized forebrain organoids, we designed a microfluidic device that can introduce 4 different growth factors into a single chamber where an organoid is embedded in hydrogels (Fig. 1A). The microfluidic device was placed on a glass slide and consisted of 5 layers, including 3 microfluidic layers and 2 porous membranes (Fig. 1B,C). The microfluidic layers were made of polydimethylsiloxane (PDMS), which is widely used for microdevices because of its biocompatibility, gas permeability, and transparency. The molds for the microfluidic layers were fabricated using a 3D printer (Methods in ESI and Fig. S1, S2).^25,26^ The middle microfluidic layer included an organoid culture chamber (5 mm (*L*) × 5 mm (*W*) × 2.5 mm (*H*)) to accommodate a forebrain organoid. The organoid culture chamber was sandwiched by 2 porous membranes that separated the upper and lower microfluidic channels. The porous membrane allowed molecular diffusion from an upper or lower channel into the chamber. The upper or lower microfluidic layers had 2 inlets to introduce 2 kinds of growth factors from each layer, leading to up to 4 growth factor inputs.

**Fig. 1.**
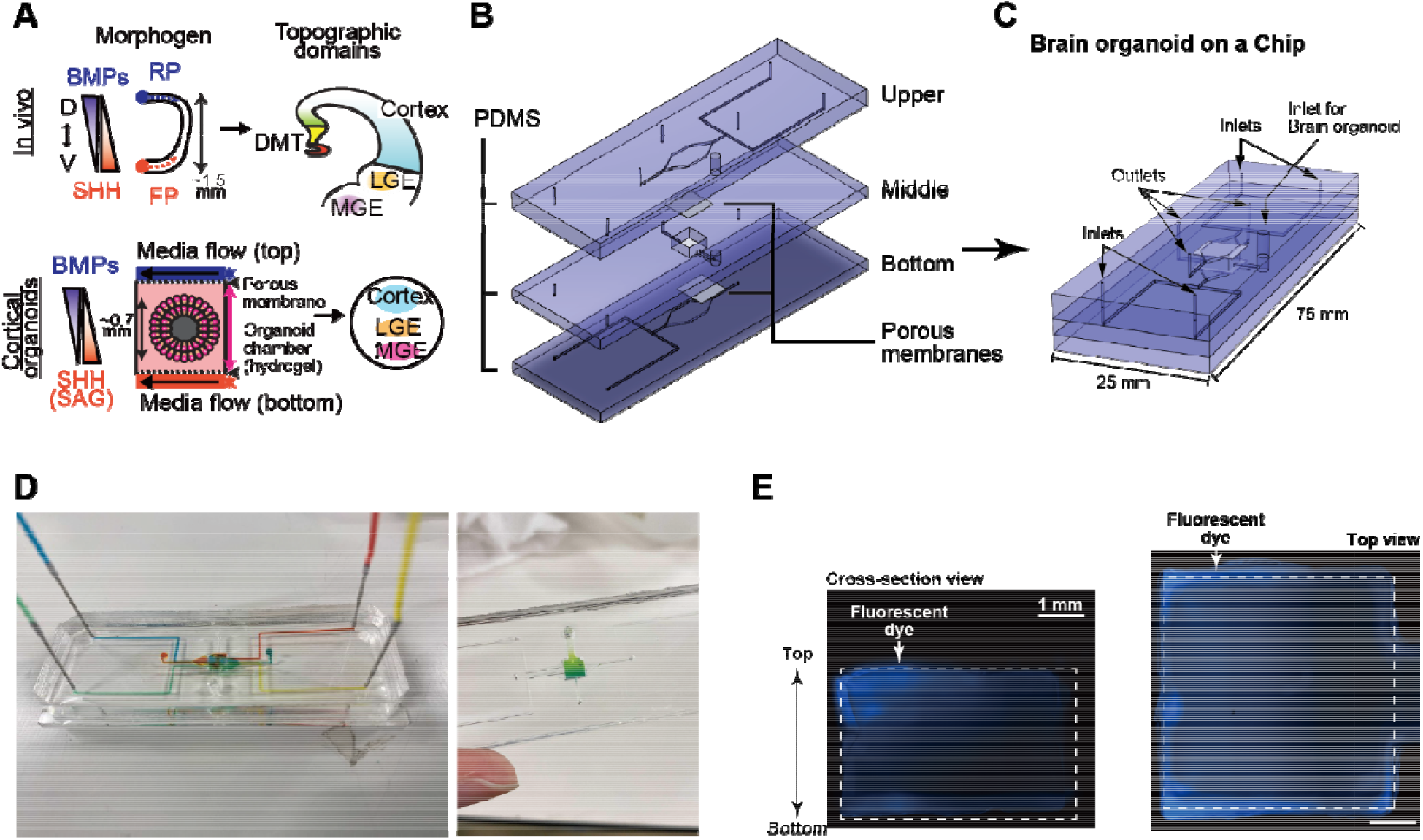
Brain organoid on a Chip. (a) Conceptual illustration of Brain-organoid-on-a-Chip (BOoC) to form a forebrain organoid with multiple domains. During forebrain development, signaling molecules (morphogen), such as BMPs and SHH, are secreted from signaling centers, the roof plate (RP) and floor plate (FR) respectively. The signaling gradients are formed to specify diverse brain regions including the dorsal midline telencephalon (DMT), cortex, lateral ganglionic eminence (LGE), and medial ganglionic eminence (MGE). (b and c) BOoC consists with 5 layers, including 3 microfluidic layers made of polydimethylsiloxane (PDMS) and 2 PET porous membranes. (d) An actual Brain organoid on a Chip introduced with colored food dyes to visualize microfluidic channels. (e) Fluorescent micrographs of agarose gel stained with fluorescent dye (AMCA-X) introduced by one of four inlets to visualize the concentration gradient in a gel. Scale bars represent 1 mm.

To maintain the location of the forebrain organoid in the chamber, thermoresponsive hydrogels [HG, a copolymer of poly(*N*-isopropyl acrylamide), and poly(ethylene glycol) (PNIPAAm-PEG)]^27–29^ were used. Since the sol-gel transition of these hydrogels can be regulated via a temperature change, the organoid can be seeded in the chamber or harvested from the chamber at a low temperature (<20 °C) and cultured in the gelated hydrogel at 37 °C. Culture medium with or without signaling molecules was diffused through the upper and lower channels through the porous membranes. In addition, the hydrogel provided a 3D structure that made it easier to generate and maintain signaling molecule concentration gradients.^26^

To confirm concentration gradients in the organoid chamber, food dyes and [6-((7-Amino-4-methylcoumarin-3-acetyl)amino) hexanoic acid] (AMCA-X) fluorescent dye were added to the medium in the device (Fig. 1D,E). The food dyes allowed visualization of perfusion flow in the chip. The dyes traversed through their individual channels and separately entered the organoid chamber via each corresponding outlet. To visualize how soluble factors diffuse across the hydrogel in the organoid chamber, a solution with AMCA-X dye was introduced into one of the inlets while the other three inlets lacked dye but were at the same flow rate (1.0 μL min^-1^). AMCA-X dye penetrated through the PET porous membrane and reached the hydrogel in the organoid chamber. Thus, the soluble factors were able to access the organoid chamber.

### Topographic forebrain organoids

We previously established a highly efficient method to generate cortical organoids from human PSCs (hPSCs) (Fig. 2A, B).^4,7^ Central nervous system induction during development requires the inhibition of Transforming Growth Factor-β (TGFβ) superfamily signaling pathways. ^30^ The default identity is the most dorsal part of the brain, the cerebral cortex, ^30,31^. Here, we began by inhibiting TGFβ and Wingless (WNT) pathways to direct hPSCs to a cortical identity. We then activated the ventralizing signal by using Sonic Hedgehog (SHH) or small molecules acting on the SHH pathway to form additional ventral brain regions, including the lateral and medial ganglionic eminence (LGE, MGE) along the dorsoventral axis (Fig. 1A). The LGE is adjacent to the cortex (Fig. 1A) and its majority becomes the striatum, important for motor and action planning, motivation, and reward perception. The MGE is further ventral adjacent to the LGE and produces the majority of GABAergic cells (Fig. 1A).^10,32^

**Fig. 2.**
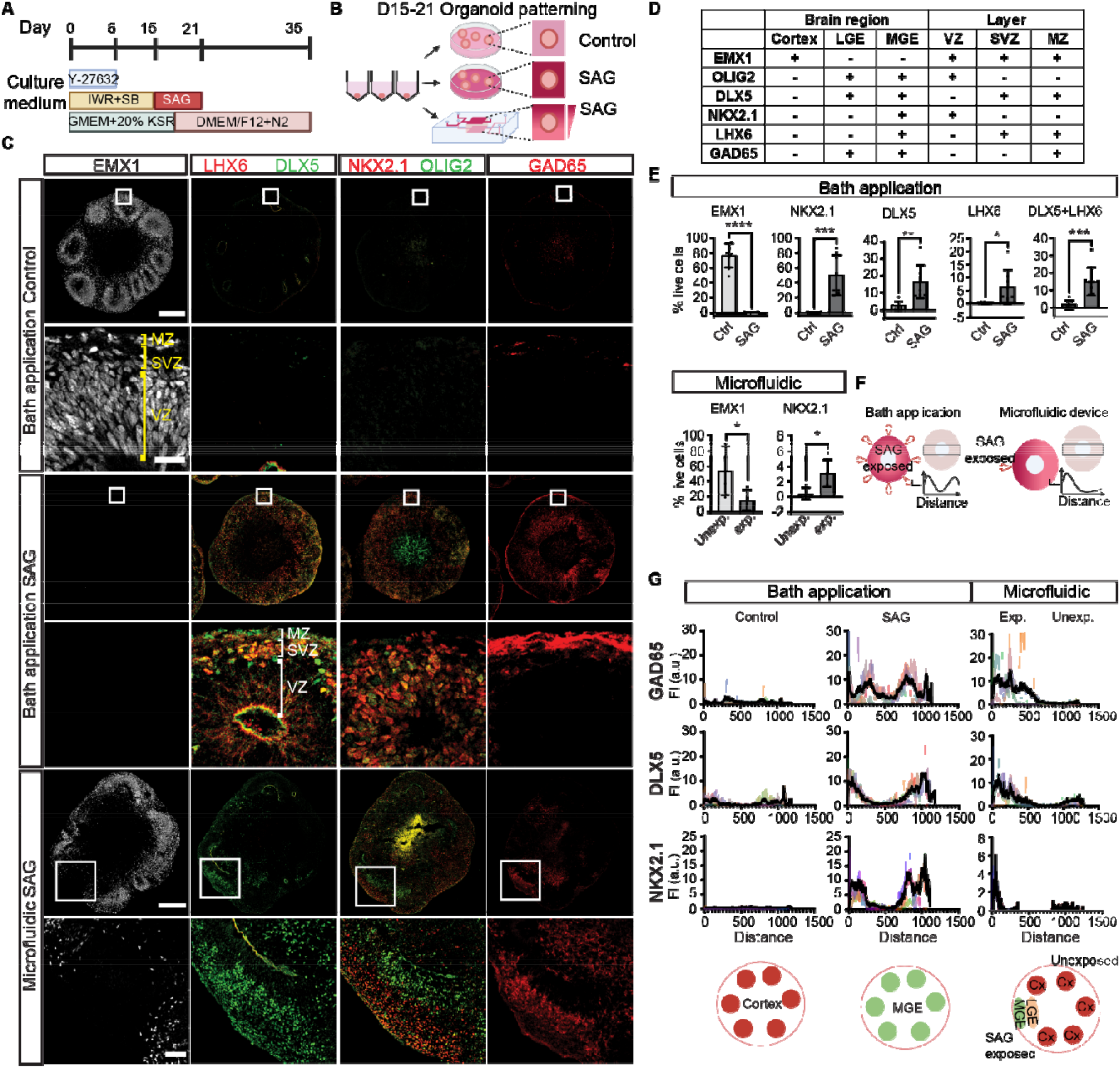
Generating topographic organization in the forebrain organoids. (a) The illustration of culture conditions for the cerebral brain organoids from day 0-35. (b) The schematic diagram of the experimental design. The SAG exposure, either with bath application or using microfluidic chips, was carried out from 15-21 days in culture. (c) Immunohistochemical analyses of forebrain organoids from control, bath SAG, and microfluidic SAG groups. Organoids from control and experimental groups were sectioned and stained for region- and cell-type-specific markers. Lower panels: zoom-in images of the box regions. MZ: mantle zone, SVZ: subventricular zone, VZ: ventricular zone. Right panels: illustrations of the regional identities of the organoids. Cortex marker: EMX1; GE markers: DLX5 and OLIG2; MGE markers: LHX6, NKX2.1; Inhibitory neuron marker: GAD65. Scale bars for bath application: 250 μm (whole), 25 μm (zoom-in). Scale bars for microfluidic chip: 250 _m (whole), 50 μm (zoom-in). (d) List of cell-type- and region-specific markers used in this study and their expression patterns. (e) Percentage of live cells with region-specific markers in organoids from bath application control (n=8), bath SAG (n=8), and microfluidic chips (n=4). Unexp: unexposed side, exp: exposed side. Bath application statistics: unpaired t-Test. Microfluidic chip statistics: paired t-Test. *p≤0.05, **p≤0.01, ***p≤0.001. (f) Illustrations of topographic quantifications of organoids from bath application groups and microfluidic chips. Fluorescent intensity is measured across the organoids from edge to edge. Height of the box: 250 μm. (g) Topographic quantifications of cell-type- and region-specific markers in organoids from bath application control (n=5), SAG (n=5), and microfluidic SAG (n=4, n=2 showed NKX2.1 expression). Illustrations (bottom panels) show the regional identities of the organoids.FI: fluorescent intensity, a.u.: arbitrary unit.

Briefly, we dissociated hPSCs into single cells and plated 9000 cells/well into a V-shaped 96-well plate. After a couple of hours, the cells coalesced to form a relatively uniform aggregate per well. We inhibited WNT and TGFβ signaling pathways, and the aggregates self-organized into early forebrain neuroepithelial cells positive for EMX1 (∼80%) at 35 days *in vitro* with some cortical lamination (Fig. 2C,D).

By default, after TGFβ inhibition, forebrain organoids become the dorsal structure, cortex. To induce ventral forebrain, the ganglionic eminence (Fig 1A), we first determined the concentrations of SAG needed to reliably form LGE and MGE. With bath application (directly adding the molecules to the culture medium), the SHH signaling pathway was activated for six days from D15 to D21 and characterized organoids at D35 (Fig. 2A,B, S2C). With bath application of 250 nM SAG, the expression of a cortical progenitor marker, EMX1, was decreased (Fig. S2C), whereas other markers such as DLX5, OLIG2, and GAD65, were upregulated, suggesting a partial LGE identity (Fig. S2C). Application of 500 nM SAG greatly diminished EMX1-positive cells, showing a loss in dorsal identity, and markers for general GE (OLIG2, DLX5, and GAD65) and MGE (NKX2.1 and LHX6) were upregulated, indicating increased ventral characteristics (Fig. 2C, S2C). Collectively, we found distinct concentrations of SAG that can induce cortical to LGE regions (250 nM) and MGE regions (500 nM) in an organoid with bath applications. Similarly, SHH gave a concentration-dependent upregulation of ventral markers (Fig. S2D). Together, SHH activation enabled the induction of LGE and MGE regions in a dose-dependent manner with bath application of signaling molecules. These data suggest that a gradient of similar molecules may aid the generation of dorsal and ventral regions in a single organoid.

Next, we tested whether the application of a signaling gradient would affect the formation of dorsal and ventral domains. We loaded neuroectodermal organoids and the hydrogel solution into the organoid chamber of the microfluidic device on ice and then warmed to 37°C for gelation (Fig. 2B). The hydrogel-embedded organoid was exposed to SAG from the upper channel from D15 to D21, creating a gradient from 500 nM SAG (Fig. 2A,B). At D21, we removed the organoid from the chamber and continued culture in a well plate in neural maintenance media until D35. We found the SAG-exposed side of the organoid had ventral character (positive for LGE and/or MGE markers and negative for cortical markers) (Fig. 2C). In contrast, the other side retained dorsal characteristics (positive for cortical markers but negative for LGE/MGE markers) (Fig. 2C).

A comparison of the organoids exposed to the SAG bath application to those exposed to the SAG gradient revealed distinct domain patterns. Organoids exposed to bath application of 250nM SAG or 10 ng/ml SHH had both dorsal and ventral regions, but they formed stochastically in space (Fig. 2C,E, S2C,D). Similarly, bath application of 50 ng/ml SHH induced both LGE and MGE regions stochastically (Fig. S2D). In contrast, regions of organoids exposed to higher concentrations of SAG (500-1000 nM) in the microfluidic gradient formed ventral domains (LGE and/or MGE markers), whereas those experiencing low concentrations generated dorsal domains (cortex markers) (Fig. 2C,E,G). This precise spatial organization of ventral and dorsal domains was not consistently observed with bath application. Together, our multilayered microfluidic device enabled a gradient of SAG exposure to induce spatially organized dorsal and ventral regions.

## Discussion

The design of our multilayered microfluidic device enabled the generation of four signaling gradients into an organoid chamber. Using gradients in the device, we formed multi-region brain organoids by ventralizing one side of the organoid. Bath application of the signaling molecule SAG induced ventral structures, but they were not spatially organized. In contrast, exposure to a SAG gradient in the microfluidic device induced dorsal and ventral regions in a spatially organized manner. Our results demonstrate a reproducible and well-controlled method for generating topographically organized brain organoids with dorsoventral structures.

Developing multi-region forebrain organoids is of great interest for accurately studying human neurodevelopment and disease. Different brain regions generate region-specific cell types for distinct functions. For example, inhibitory interneurons are generated mainly in ventral structures, specifically the MGE and caudal ganglionic eminence (CGE), ^33^ and migrate tangentially toward the cerebral cortex. Interneurons comprise about 20-30% of neurons in the cortex and interact with excitatory glutamatergic neurons in the cerebral cortex to regulate neuronal circuitry.^34^ If ventral MGE/CGE regions are not induced in cortical organoids, the lack of inhibitory interneurons means that proper neural microcircuits cannot form. Another approach to generating regionalized organoids is to fuse organoids of different regional identities together, creating assembloids. Fusing cortical and GE organoids increased spontaneous neuronal activity and led to more complex neuronal oscillations.^11^ However, cell migration from ventral to dorsal structures is a critical hallmark of brain development and is lacking in assembloids. Also, region-region communication begins at early time points, so it is desirable for brain regions to develop together, ideally in a spatially organized manner to mimic physiological conditions. Our microfluidic device can form multiple gradients of signaling molecules, raising the possibility of generating more than two brain regions simultaneously in a single organoid. This is important for studying neurological and psychiatric disorders using human brain organoids, as multiple brain regions and different cell types may be involved in pathophysiology.

Key factors in the developing brain, including BMPs, FGFs, and SHH, are secreted from signaling centers, and the concentration, timing, and spatial distribution of these signaling molecules determine cell fate.^35^ For example, SHH is initially secreted from the notochord and the floor plate for ventral patterning and later from the *zona limitans intrathalamica* for oligodendrocyte specification.^36^ The failure to form signaling centers during forebrain development leads to holoprosencephaly, the most common malformation of the human forebrain.^37^ Our multilayered microfluidic device supplied SAG on one side of the organoid chamber, mimicking the action of the ventral signaling center and enabling the generation of spatially organized ventral structures in forebrain organoids. Whether the induction of ventral cell fate using the device is sufficient to induce the formation of the signaling center is yet to be investigated.

An advantage of the device is the ability to supply multiple signaling molecules on each side of the organoid. Further, the design provides flexibility for the user in controlling the concentration and timing of a given signaling molecule. A limitation of this approach is the use of perfusion flow to generate signaling molecule concentration gradients. While our device design prevents the direct application of shear stress to the organoid, the perfusion flow used to create the gradient leads to significant consumption of signaling molecules. Since many signaling molecules are quite expensive, cost considerations limit the duration of experiments. To overcome this limitation, we employed the less-expensive SAG as a substitute for SHH. However, their diffusion coefficients differ, and the use of SAG might not fully replicate the in vivo developmental process. Future work should explore alternative methods to perfusion flow for applying concentration gradients to minimize the consumption of signaling molecules.

The production of topographically organized organoids is low throughput because the current design is limited to only one organoid per chip. Future designs will accommodate multiple organoids in each chip for higher throughput. The current design of the organoid chamber does not control the landing position of the organoid, which can affect the part of the concentration gradient experienced by each organoid. More refined control of the organoid position is needed to further reduce variability in topographically organized forebrain organoid formation.

## Conclusions

In this study, we used a multilayered microfluidic device to mimic the diffusion of molecules from brain signaling centers to guide the differentiation of human organoids. With this method, we successfully generated organoids comprised of dorsal (cortical) and ventral (LGE and MGE) regions in a spatially organized manner, mimicking the brain development along the dorsoventral axis. This system is applicable to different types of organoids and can be utilized for drug delivery.

## Author Contributions

MW and KK conceptualized the study. KK, AM and SMH designed and fabricated multilayered microfluidic devices. YCT and HO performed experiments on forebrain organoids. KS generated and maintained forebrain organoids. APP, AGS, KAP, BKS, and CHV quantified organoid datasets. YCT performed statistical analyses. MW, KK, and YCT wrote the manuscript.

## Acknowledgements

Funding: This work was supported by the NIH R00HD096105, NSF RECODE2225624, New Investigator Faculty Award and start-up funds from the UCI School of Medicine to MW, the FRAXA Postdoctoral Fellowship to YCT, the Japan Society for the Promotion of Science (JSPS) (21H01728) and the Japan Agency for Medical Research and Development (AMED) (17937667) to KK. WPI-iCeMS is supported by the World Premier International Research Centre Initiative (WPI), MEXT, Japan. The authors acknowledge Dr. Lisa A. Flanagan for critical reading of the manuscript. Illustrations in figure 2 and supplementary figure 3 were created with BioRender.com.

## References

1 1 T. Kadoshima, H. Sakaguchi, T. Nakano, M. Soen, S. Ando, M. Eiraku and Y. Sasai, Proceedings of the National Academy of Sciences, 2013, 110, 20284–20289.

2 M. A. Lancaster, M. Renner, C. A. Martin, D. Wenzel, L. S. Bicknell, M. E. Hurles, T. Homfray, J. M. Penninger, A. P. Jackson and J. A. Knoblich, Nature, 2013, 501, 373–379.

3 M. Eiraku, K. Watanabe, M. Matsuo-Takasaki, M. Kawada, S. Yonemura, M. Matsumura, T. Wataya, A. Nishiyama, K. Muguruma and Y. Sasai, Cell Stem Cell, 2008, 3, 519–532.

4 M. Watanabe, J. E. Buth, N. Vishlaghi, L. de la Torre-Ubieta, J. Taxidis, B. S. Khakh, G. Coppola, C. A. Pearson, K. Yamauchi, D. Gong, X. Dai, R. Damoiseaux, R. Aliyari, S. Liebscher, K. Schenke-Layland, C. Caneda, E. J. Huang, Y. Zhang, G. Cheng, D. H. Geschwind, P. Golshani, R. Sun and B. G. Novitch, Cell Rep, 2017, 21, 517–532.

5 A. Bhaduri, M. G. Andrews, W. Mancia Leon, D. Jung, D. Shin, D. Allen, D. Jung, G. Schmunk, M. Haeussler, J. Salma, A. A. Pollen, T. J. Nowakowski and A. R. Kriegstein, Nature 2020 578:7793, 2020, 578, 142–148.

6 S. J. Yoon, L. S. Elahi, A. M. Pa_ca, R. M. Marton, A. Gordon, O. Revah, Y. Miura, E. M. Walczak, G. M. Holdgate, H. C. Fan, J. R. Huguenard, D. H. Geschwind and S. P. Pa_ca, Nat Methods, 2019, 16, 75–78.

7 M. Watanabe, J. E. Buth, J. R. Haney, N. Vishlaghi, F. Turcios, L. S. Elahi, W. Gu, C. Pearson, A. Kurdian, N. V. Baliaouri, A. J. Collier, O. A. Miranda, N. Dunn, D. Chen, S. Sabri, L. de la Torre-Ubieta, A. T. Clark, K. Plath, H. R. Christofk, H. I. Kornblum, M. J. Gandal and B. G. Novitch, Stem Cell Reports, 2022, 17, 2220–2238.

8 S. Velasco, A. J. Kedaigle, S. K. Simmons, A. Nash, M. Rocha, G. Quadrato, B. Paulsen, L. Nguyen, X. Adiconis, A. Regev, J. Z. Levin and P. Arlotta, Nature, 2019, 570, 523–527.

9 J. A. Bagley, D. Reumann, S. Bian, J. Lévi-Strauss and J. A. Knoblich, Nat Methods, 2017, 14, 743–751.

10 F. Birey, J. Andersen, C. D. Makinson, S. Islam, W. Wei, N. Huber, H. C. Fan, K. R. C. Metzler, G. Panagiotakos, N. Thom, N. A. O’Rourke, L. M. Steinmetz, J. A. Bernstein, J. Hallmayer, J. R. Huguenard and S. P. Pasca, Nature, 2017, 545, 54–59.

11 R. A. Samarasinghe, O. A. Miranda, J. E. Buth, S. Mitchell, I. Ferando, M. Watanabe, T. F. Allison, A. Kurdian, N. N. Fotion, M. J. Gandal, P. Golshani, K. Plath, W. E. Lowry, J. M. Parent, I. Mody and B. G. Novitch, Nature Neuroscience 2021 24:10, 2021, 24, 1488–1500.

12 Y. Xiang, Y. Tanaka, B. Patterson, Y. J. Kang, G. Govindaiah, N. Roselaar, B. Cakir, K. Y. Kim, A. P. Lombroso, S. M. Hwang, M. Zhong, E. G. Stanley, A. G. Elefanty, J. R. Naegele, S. H. Lee, S. M. Weissman and I. H. Park, Cell Stem Cell, 2017, 21, 383-398.e7.

13 Y. Xiang, Y. Tanaka, B. Cakir, B. Patterson, K. Y. Kim, P. Sun, Y. J. Kang, M. Zhong, X. Liu, P. Patra, S. H. Lee, S. M. Weissman and I. H. Park, Cell Stem Cell, 2019, 24, 487-497.e7.

14 M. A. Lancaster, M. Renner, C. A. Martin, D. Wenzel, L. S. Bicknell, M. E. Hurles, T. Homfray, J. M. Penninger, A. P. Jackson and J. A. Knoblich, Nature, 2013, 501, 373–379.

15 A. G. Monteduro, S. Rizzato, G. Caragnano, A. Trapani, G. Giannelli and G. Maruccio, Biosens Bioelectron,, DOI:10.1016/j.bios.2023.115271.

16 D. E. Ingber, Nature Reviews Genetics 2022 23:8, 2022, 23, 467–491.

17 Z. Ao, H. Cai, D. J. Havert, Z. Wu, Z. Gong, J. M. Beggs, K. Mackie and F. Guo, Anal Chem, 2020, 92, 4630–4638.

18 Z. Ao, S. Song, C. Tian, H. Cai, X. Li, Y. Miao, Z. Wu, J. Krzesniak, B. Ning, M. Gu, L. P. Lee and F. Guo, Advanced Science, 2022, 9, 2200475.

19 Y. Wang, L. Wang, Y. Zhu and J. Qin, Lab Chip, 2018, 18, 851–860.

20 X. Qian, H. N. Nguyen, M. M. Song, C. Hadiono, S. C. Ogden, C. Hammack, B. Yao, G. R. Hamersky, F. Jacob, C. Zhong, K. J. Yoon, W. Jeang, L. Lin, Y. Li, J. Thakor, D. A. Berg, C. Zhang, E. Kang, M. Chickering, D. Nauen, C. Y. Ho, Z. Wen, K. M. Christian, P. Y. Shi, B. J. Maher, H. Wu, P. Jin, H. Tang, H. Song and G. L. Ming, Cell, 2016, 165, 1238–1254.

21 Y. Wang, L. Wang, Y. Zhu and J. Qin, Lab Chip, 2018, 18, 851–860.

22 B. Cakir, Y. Xiang, Y. Tanaka, M. H. Kural, M. Parent, Y. J. Kang, K. Chapeton, B. Patterson, Y. Yuan, C. S. He, M. S. B. Raredon, J. Dengelegi, K. Y. Kim, P. Sun, M. Zhong, S. Lee, P. Patra, F. Hyder, L. E. Niklason, S. H. Lee, Y. S. Yoon and I. H. Park, Nature Methods 2019 16:11, 2019, 16, 1169–1175.

23 M. A. Qasaimeh, T. Gervais and D. Juncker, Nature Communications 2011 2:1, 2011, 2, 1–8.

24 24 W. Lim and S. Park, Molecules 2018, Vol. 23, Page 3355, 2018, 23, 3355.

25 R. Abdalkader and K. Kamei, Lab Chip, 2020, 20, 1410–1417.

26 K. Kamei, Y. Mashimo, Y. Koyama, C. Fockenberg, M. Nakashima, M. Nakajima, J. Li and Y. Chen, Biomed Microdevices, 2015, 17, 36.

27 H. Yoshioka, Y. Mori, S. Tsukikawa and S. Kubota, Polym Adv Technol, 1998, 9, 155–158.

28 H. Yoshioka, Y. Mori and J. A. Cushman, Polym Adv Technol, 1994, 5, 122–127.

29 K. I. Kamei, Y. Koyama, Y. Tokunaga, Y. Mashimo, M. Yoshioka, C. Fockenberg, R. Mosbergen, O. Korn, C. Wells and Y. Chen, Adv Healthc Mater, 2016, 5, 2951–2958.

30 T. Kadoshima, H. Sakaguchi, T. Nakano, M. Soen, S. Ando, M. Eiraku and Y. Sasai, Proc Natl Acad Sci U S A, 2013, 110, 20284–20289.

31 M. Eiraku, K. Watanabe, M. Matsuo-Takasaki, M. Kawada, S. Yonemura, M. Matsumura, T. Wataya, A. Nishiyama, K. Muguruma and Y. Sasai, Cell Stem Cell, 2008, 3, 519–532.

32 Y. Miura, M. Y. Li, F. Birey, K. Ikeda, O. Revah, M. V. Thete, J. Y. Park, A. Puno, S. H. Lee, M. H. Porteus and S. P. Pa□ca, Nature Biotechnology 2020 38:12, 2020, 38, 1421–1430.

33 D. V. Hansen, J. H. Lui, P. Flandin, K. Yoshikawa, J. L. Rubenstein, A. Alvarez-Buylla and A. R. Kriegstein, Nature Neuroscience 2013 16:11, 2013, 16, 1576–1587.

34 H. Markram, M. Toledo-Rodriguez, Y. Wang, A. Gupta, G. Silberberg and C. Wu, Nature Reviews Neuroscience 2004 5:10, 2004, 5, 793–807.

35 W. Wurst, L. Bally-Cuif and L. Bally-Cuif, Nature Reviews Neuroscience 2001 2:2, 2001, 2, 99–108.

36 E. Martí and P. Bovolenta, Trends Neurosci, 2002, 25, 89–96.

37 E. S. Monuki, J Neuropathol Exp Neurol, 2007, 66, 566–575.

