## Supplementary information for "Brain organoid-on-a-chip to create multiple domains in forebrain organoids"

**Electronic supplementary information for**

**Material and Methods**

**1. Microfluidic device fabrication**

Microfluidic devices were fabricated using stereolithographic 3D-printing techniques and molding processes (Fig. S1).[1,2] The molds for the top, middle and bottom layers were produced using a 3D printer (Keyence Corporation, Osaka, Japan). After printing, the molds were washed with 99.9% EtOH at 25°C for 12 h, and dried at 80 °C for 30 min. The microfluidic channels are designed with 100 µm in width and 500 µm in height, and have 0.4-mm inlets. Before use, the surfaces of the molds were coated with trichloro(1H,1H,2H,2H-perfluorooctyl) silane (Sigma-Aldrich, St. Louis, MO, USA). 33 g of polydimethylsiloxane (PDMS; Sylgard 184 PDMS two-part elastomers, Dow Corning Corporation, Midland, MI, USA) was mixed with 10:1 ratio of pre-polymer and curing agent, and poured into mold to produce 4-mm, 2.5-mm and 2.5-mm thick PDMS upper, middle and lower layers, respectively, and de-gassed using a vacuum desiccator at 25°C for 30 min. All molds with pre-PDMS mixer were cured in an oven at 80°C for 20 min. After curing, solidified PDMS structures were peeled off from the molds, trimmed, and cleaned. The upper and middle layers were peeled off from the molds, and treated with a corona plasma (Shinko Denki, Inc., Oosaka, Japan). Then, a PET membrane (pore size 3.0 µm; pore density $1.6 \times 10^6$ $cm^{-2}$; it4ip, Louvain-la-Neuve, Belgium) was placed between the upper and middle layers. Then, the assembled upper-middle layers and lower layer were treated with a corona plasma and assembled with a PET membrane. The lower layer and a glass slide were treated with a corona plasma, and assembled together, followed by curing it in an oven at 80°C for 24 hr.

**2. Human embryonic stem cell (hESC) culture and organoid differentiation**

hESC experiments were conducted with prior approval from the University of California, Irvine (UCI) Human Stem Cell Research Oversight (hSCRO) Committee. hESC line H9 was obtained from the WiCell Institute after the material transfer agreement. For the quality control, we checked chromosomal abnormality using hPSC Genetic Analysis Kit (Stemcell Technologies) and mycoplasma infection using Mycoplasma Detection Kit (SouthernBiotech™) every 10 passages.

H9 cells between passage 50 to 80 were seeded with inactivated mouse embryonic feeders (PMEF-CF, Millipore Sigma) on 0.1% gelatin-coated plate, and cultured in DMEM/F12 (HyClone) with 20% knockout serum replacement (KSR, Invitrogen), non-essential amino acids (NEAAs, Invitrogen), GlutaMAX (Invitrogen), 100 mg/mL of Primocin (InvivoGen), 0.1 mM beta-mercaptoethanol (Invitrogen), and 10 ng/mL of fibroblast growth factor 2 (FGF2, Invitrogen). hESCs were incubated at 5% $CO_2$ at 37°C with daily media change and were passaged every 6 days at 1:3-1:7 using StemPro EZ Passage tool (Invitrogen).

The organoids were generated from hESCs as previously established [3,4]. Briefly, when hESCs were about ~70-80% confluent, hESCs were dissociated to single cells and plated into low-attachment V-bottom 96-well plates (9000 cells/well; Sumitomo Bakelite, MS9096V) to form an aggregate/well. The neural differentiation media for day 0 (D0) to D15 organoids consisted of Glasgow's Minimal Essential Medium (GMEM, Invitrogen), 20% KSR, NEAA, 100 mg/mL of Primocin, 0.1 mM b-mercaptoethanol, sodium pyruvate (Invitrogen), Wnt inhibitor IWR-1-endo (Calbiochem), and TGF-b inhibitor SB431542 (Stemgent). 20 μM ROCK inhibitor Y-27632 (BioPioneer) was also supplied in the medium from D0 to D6. Half of the culture media was changed every 2-3 days from D6 to D15. D0-D15 organoids were maintained at 5% $CO_2$ at 37°C.

**3. Organoid ventralization**

The organoids were maintained at 5% $CO_2$ and 40% $O_2$ at 37°C from D15 to D35. The D15 organoids were subject to either bath application of varying concentrations of SAG (250-750 nM) or embedded in an organoid chamber of the microfluidic device, where SAG (500-1000 nM) media was applied through the inlet channels.

3.1 Bath application

The D15 organoids were transferred to Petri dishes and bathed with SAG in the neural differentiation medium, and the media with SAG were refreshed on D18. From D21, the media were changed to the neural maintenance media, including DMEM/F12 with N2 (Invitrogen), GlutaMAX, chemically defined lipid concentrate (CDLC, Invitrogen), and 0.4% methylcellulose (Sigma). The culture media was changed every 2–3 days and the organoids were maintained until D35.

#### 3.2 Microfluidic device

A D15 organoid was embedded into the MeBiol thermoreversible hydrogel (Advanced BioMatrix) and loaded into the organoid chamber of the microfluidic chip. The neural differentiation media with or without 500-1000 nM of SAG were loaded into 10 mL syringes. The syringes were placed on syringe pumps (Legato) and the PTFE tubing connected the syringes to the microfluidic chip. The neural differentiation media without SAG was supplied through the lower channels, and the neural differentiation media with SAG 500nM was supplied through the upper channels of the microfluidic chip. The injecting speed was set to 1 μL per minute and the organoid was incubated in the microfluidic chip for 6 days. On D21, the organoid was removed from the microfluidic chip and transferred to a well of the 24-well plate. From D21, the neural differentiation media was changed to the neural maintenance media. The culture media was changed every 2–3 days and the organoids were maintained until D35.

### 4. Immunohistochemistry

At D35, organoids were fixed in 4% paraformaldehyde/PBS solution, cryprotected in 30% sucrose/PBS solution, and embedded in Tissue-Tek Optimal Cutting Temperature medium (Sakura, 25608-930). Samples were cryosectioned at 12 μm. Immunostaining was performed as previously described[3–5]. In short, organoid cryosections were incubated in blocking solution (0.1% heat-inactivated horse serum, 0.1% Triton-X, 0.01% sodium azide, PBS) for one hour at room temperature prior to primary antibody incubation at 4°C overnight. After washing with PBST (0.1% TritonX), the tissue sections were then incubated in secondary antibody solutions, including Hoechst 33258 for nuclei staining, for one hour at room temperature. After washing with PBST, the cryosections were mounted in Prolong Diamond Antifade, covered with coverslips, and subjected to confocal imaging. Primary antibodies used in this study were the following: guinea pig anti-DLX5 (BioAcademia, 74-117) 1:2000 dilution; rabbit anti-EMX1 (Sigma-Aldrich, HPA006421) 1:100 dilution; mouse anti-GAD65 (BD Biosciences, 559931) 1:200 dilution; mouse anti-LHX6 (Santa Cruz, sc-271433) 1:200 dilution; mouse anti-NKX2.1 (Novocastra, NCL-L-TTF-1) 1:500 dilution, rabbit anti-OLIG2 (Millipore, AB9610) 1:5000 dilution, goat anti-SOX2 (R&D Systems, AF2018) 1:1000 dilution. Secondary antibodies used in this study are the following: Alexa 647-, Alexa 568- , Alexa 488-Donkey anti-species-specific IgG (Jackson ImmunoResearch) at 1:700 dilution for Alexa 647 and 1:500 dilution for Alexa 568 and Alexa 488.

For cell counting, the number of EMX1+, NKX2.1+, DLX5+, and LHX6+ cells are counted out of total live cells, using ImageJ/Fiji, and the percentages were calculated. The necrotic core of the organoids was excluded from the cell counting to only account for live cells. For topographic analysis, images were rotated so that the SAG exposure side was on the left side. The levels of fluorescent intensity were measured across the organoids (length = the length of the organoids, width = 250 µm) using ImageJ/Fiji. The graphs were generated using the GaphPad Prism 9 software.

**5. Confocal microscopy**

Tiled images were taken with 10X objective using Zeiss LSM 900 Airy Scan 2 (Zen Blue 3.3 software). Within the same experimental sets, the images were obtained with the same settings for laser and acquisition power. ImageJ/Fiji was used for further adjustment of brightness, contrast and levels. The uniform and same image adjustments were used in the control and the experiment groups in a set. ImageJ/Fiji was used for manual cell counting on the processed images.

**6. RNA isolation and RT-qPCR**

Organoid samples were lysed with Buffer RLT (Qiagen) and total RNA was extracted using RNeasy Mini Kit (Qiagen) following the manufacturer's user manual. More than 1000 ng of RNA from each sample was used for cDNA synthesis using SuperScript IV First-Strand Synthesis Reaction (Invitrogen). For RT-qPCR, the cDNA samples were combined with PowerTrack SYBR Green Master Mix (Applied Biosystems) and primer pairs of the genes of interest (Integrated DNA Technologies). The samples were loaded into 384-well plates (Invitrogen) in duplicates and proceeded with QuantStudio 7 Real-Time PCR System (Applied Biosystems). The relative expression level of each gene was calculated by normalizing it to the housekeeping gene GAPDH. The exon-spanning primers used are the following: GAPDH (amplicon size 69 bp) fw 5'-CTCTCTGCTCCTCCTGTTCGAC-3' rv 5'-TGAGCGATGTGGCTCGGCT-3'; NKX2.1 (amplicon size 68 bp) fw 5'-AGCACACGACTCCGTTCTC-3' rv 5'-GCCCACTTTCTTGTAGCTTTCC-3'; OLIG2 (amplicon size 90 bp) fw 5'-ATAGATCGACGCGACACCAG-3' rv 5'-ACCCGAAAATCTGGATGCGA-3.

**7. Statistical information**

Unpaired Student's two-tailed t-tests and paired t-tests were performed using GaphPad Prism 9 software. The p-value and n numbers are reported in the figure legends. Significance was assumed when $p < 0.05$. Statistical significance is presented as the following: * $p \leq 0.05$, ** $p \leq 0.01$, *** $p \leq 0.001$, **** $p \leq 0.0001$. Error bar: standard deviation.

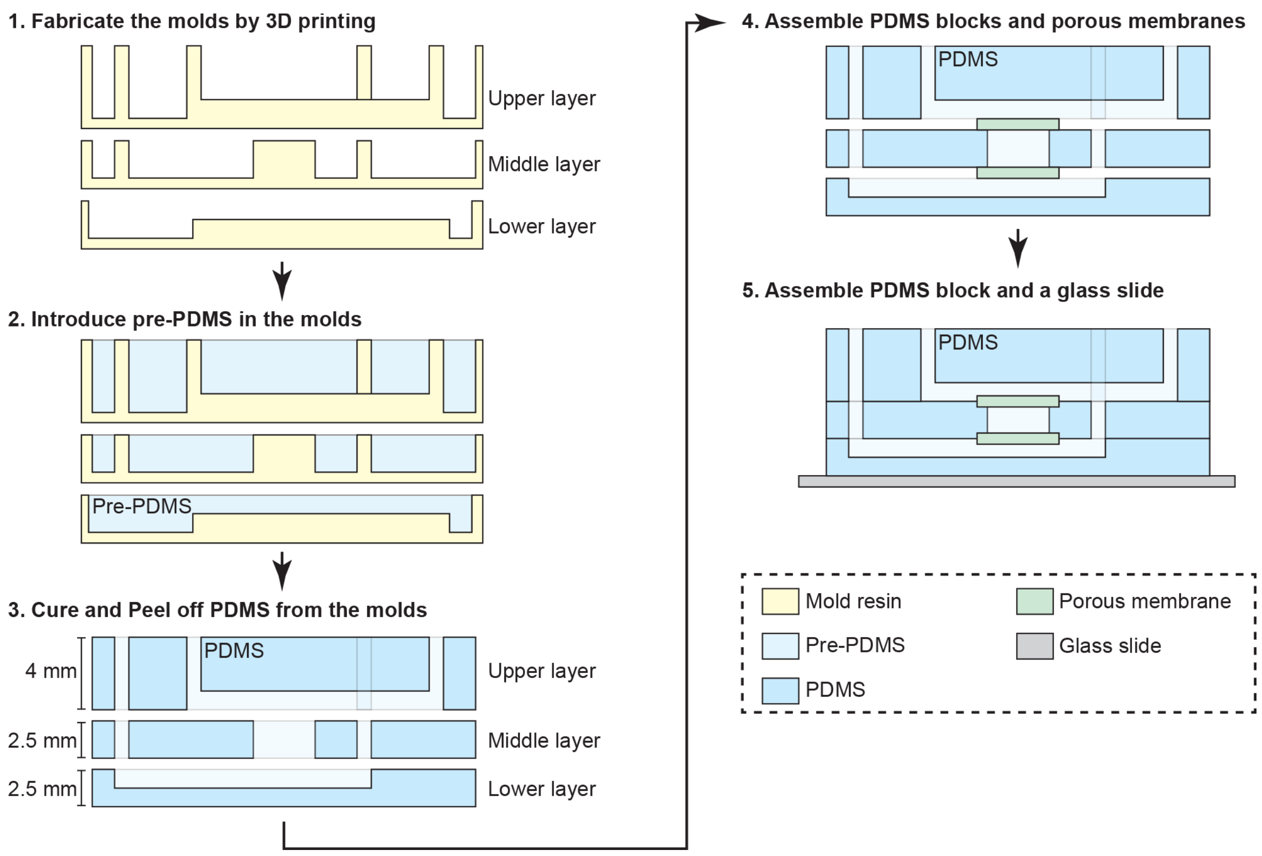


**Fig. S1.** Fabrication procedure of a microfluidic device used for Brain-on-a-Chip.

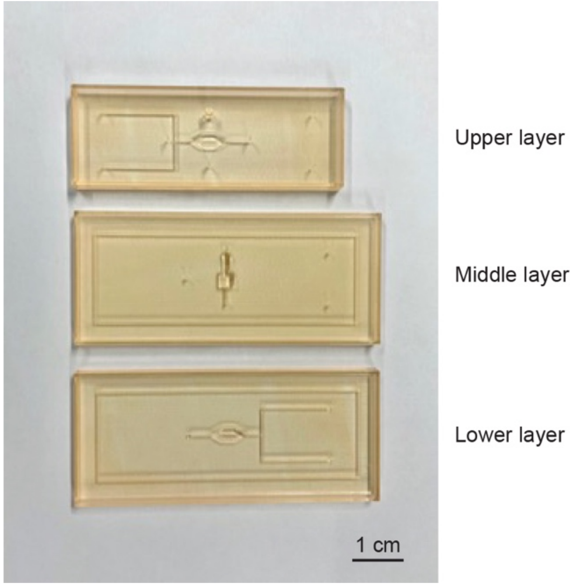


**Fig. S2** 3D-printed molds for a microfluidic device used for Brain-on-a-Chip

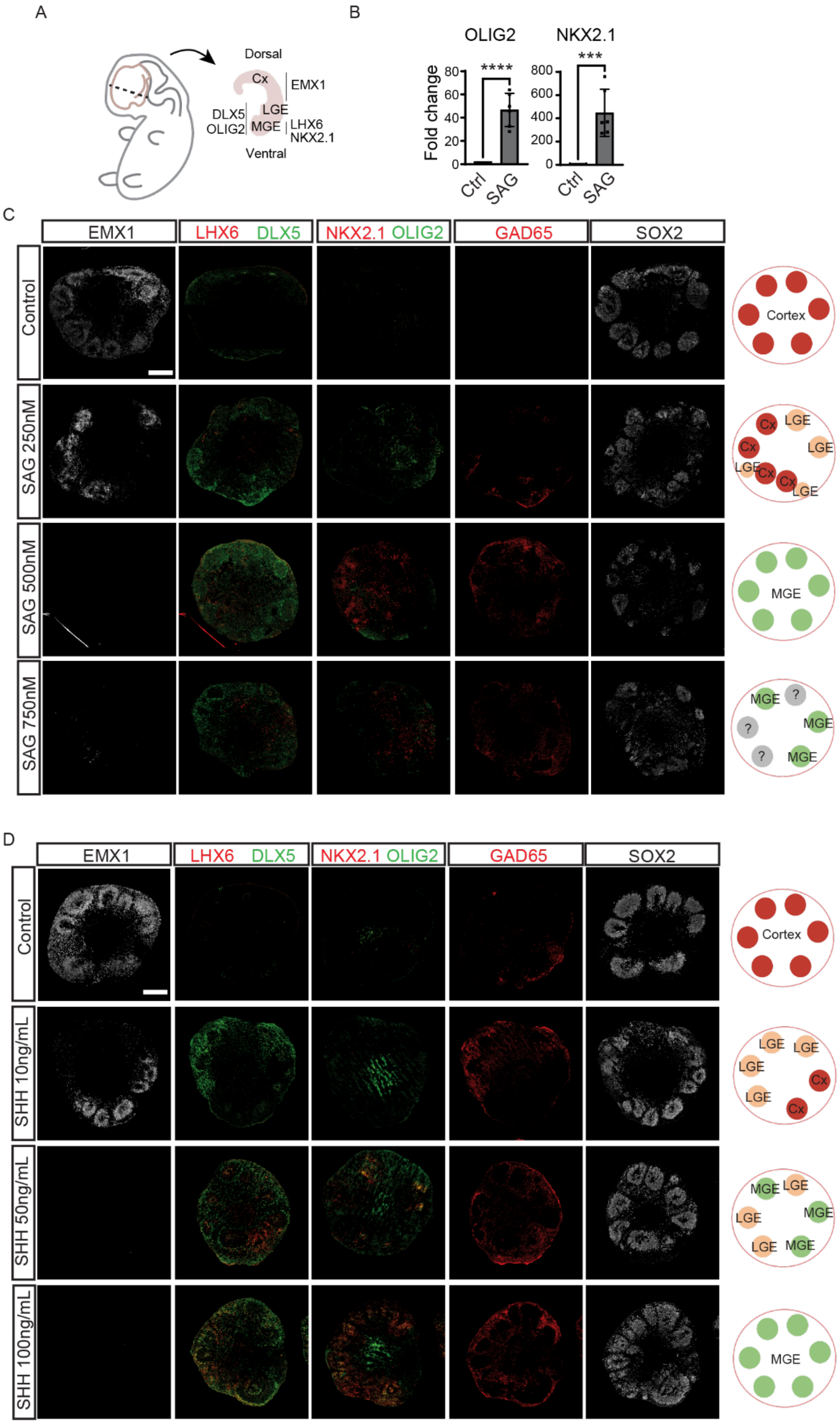
A
Dorsal
Cx
EMX1
DLX5
OLIG2
LGE
MGE
LHX6
NKX2.1
Ventral
B
OLIG2
NKX2.1
Fold change
****
***
0
20
40
60
80
0
200
400
600
800
Ctrl
SAG
Ctrl
SAG
C
EMX1
LHX6 DLX5
NKX2.1 OLIG2
GAD65
SOX2
Control
SAG 250nM
SAG 500nM
SAG 750nM
Cortex
Cx
LGE
MGE
?
D
EMX1
LHX6 DLX5
NKX2.1 OLIG2
GAD65
SOX2
Control
SHH 10ng/mL
SHH 50ng/mL
SHH 100ng/mL
Cortex
LGE
Cx
MGE

**Fig. S3** Bath application of various concentrations of SAG and SHH. (A) Illustration of the brain regions in the embryonic brain and specific markers for cortex (Cx), lateral ganglionic eminence (LGE), and medial ganglionic eminence (MGE). (B) Gene expression level of GE-specific markers in bath application control and SAG (n=6 per group). Statistics: unpaired t-Test. $^{*}p\leq0.05$, $^{**}p\leq0.01$, $^{***}p\leq0.001$, $^{****}p\leq0.0001$. (C) Shift of region identity, from cortex to GE, with increasing SAG concentrations. Images of organoids immunostained with the region- and cell-type-specific markers. Right panel: illustrations of the regional identities present in the organoids. (D) Shift of region identity, from cortex to GE, with increasing concentrations of sonic hedgehog (SHH).

### References


1 K. Kamei, Y. Mashimo, Y. Koyama, C. Fockenberg, M. Nakashima, M. Nakajima, J. Li and Y. Chen, *Biomed Microdevices*, 2015, **17**, 36.

2 R. Abdalkader and K. Kamei, *Lab Chip*, 2020, **20**, 1410–1417.

3 M. Watanabe, J. E. Buth, N. Vishlaghi, L. de la Torre-Ubieta, J. Taxidis, B. S. Khakh, G. Coppola, C. A. Pearson, K. Yamauchi, D. Gong, X. Dai, R. Damoiseaux, R. Aliyari, S. Liebscher, K. Schenke-Layland, C. Caneda, E. J. Huang, Y. Zhang, G. Cheng, D. H. Geschwind, P. Golshani, R. Sun and B. G. Novitch, *Cell Rep*, 2017, **21**, 517–532.

4 M. Watanabe, J. E. Buth, J. R. Haney, N. Vishlaghi, F. Turcios, L. S. Elahi, W. Gu, C. A. Pearson, A. Kurdian, N. V. Baliaouri, A. J. Collier, O. A. Miranda, N. Dunn, D. Chen, S. Sabri, L. de la Torre-Ubieta, A. T. Clark, K. Plath, H. R. Christofk, H. I. Kornblum, M. J. Gandal and B. G. Novitch, *Stem Cell Reports*, 2022, **17**, 2220–2238.

5 R. A. Samarasinghe, O. A. Miranda, J. E. Buth, S. Mitchell, I. Ferando, M. Watanabe, T. F. Allison, A. Kurdian, N. N. Fotion, M. J. Gandal, P. Golshani, K. Plath, W. E. Lowry, J. M. Parent, I. Mody and B. G. Novitch, *Nature Neuroscience 2021 24:10*, 2021, **24**, 1488–1500.
